# Contrasting plasmid–host associations shape the distribution of enterotoxigenic *E. coli* (ETEC) colonization factors

**DOI:** 10.64898/2026.11.16.767129

**Authors:** Biel Garcias, Marlene Grimmer, Daria Frolova, Zamin Iqbal, Sam Dougan, Enrique Joffre, Jeslin Varghese, Rhiannon Evans, David Baker, Thanh Le Viet, Andrew Page, Astrid von Mentzer

## Abstract

Enterotoxigenic *Escherichia coli* (ETEC) causes a substantial burden of diarrhoeal disease, but how its virulence plasmids spread across *E. coli* populations remains unclear. We generated 479 complete ETEC genomes and combined them with 56 publicly available assemblies to produce the largest plasmid-resolved ETEC collection to date. Clustering of 903 virulence plasmids identified distinct plasmid lineages associated with individual colonization factors (CFs). Some CFs were carried by plasmids confined to a single ETEC lineage, consistent with long-term vertical maintenance, whereas others, including CS23 and CS12, were found on closely related conjugative plasmids distributed across multiple lineages, indicating horizontal transfer. Screening the AllTheBacteria dataset confirmed these patterns and revealed additional plasmid diversity for several CFs. Finally, we identified eight previously undescribed putative adhesion operons on toxin plasmids that may explain CF-negative ETEC isolates. These findings showed that plasmids, rather than bacterial lineages alone, shape the evolution and dissemination of ETEC virulence.

## INTRODUCTION

Enterotoxigenic *Escherichia coli* (ETEC) is a major cause of diarrhoeal disease in low- and middle-income countries (LMICs), causing an estimated 220 million diarrhoeal episodes and 50,000 deaths each year^1^. Although the highest burden falls on children under five years of age^2^, ETEC is also one of the main causes of traveller’s diarrhoea^3^.

ETEC pathogenesis depends on two main steps. First, bacteria adhere to enterocytes through colonization factors (CFs). After colonization, enterotoxins induce fluid secretion into the intestinal lumen, resulting in diarrhoea^4^. Two major toxin types have been described: heat-stable toxins (ST) and heat-labile toxins (LT). In contrast, CFs are highly diverse, with 29 different operons identified so far^5^, and ETEC may harbour one or more CFs in different combinations. The most prevalent CFs are CS6, CFA/I, CS3 and CS2, which occur in different constellations such as CFA/I+CS21, CS1+CS3, CS2+CS3, CS5+CS6 or CS6 alone^6^. Additional virulence factors (VFs) with adhesive properties, such as EtpBAC^8^, have also been described, but a large proportion of ETEC strains remain without recognized CFs. This suggests that additional CFs remain to be discovered and that important aspects of ETEC epidemiology are still poorly understood.

Historically, *E. coli* pathotypes were thought to emerge simply through acquisition of VFs by any strain. This view was challenged when specific CFs were shown to be associated with conserved ETEC lineages, suggesting interactions between the core and the accessory genome that promote the long-term success of particular virulence combinations^9^. However, a recent study from Bangladesh described the emergence of CS23 in multiple phylogenetically distinct lineages, pointing to plasmid-mediated dissemination rather than simple clonal expansion^10^. Although ETEC VFs have long been known to reside on plasmids^11,12^, the role of these plasmids in shaping the distribution and spread of CFs across *E. coli* populations remains poorly understood. Most genomic surveillance studies rely on short-read sequencing, which frequently fails to accurately reconstruct plasmids because of fragmented assemblies.

To address this limitation, we combined long- and short-read sequencing to fully reconstruct ETEC genomes and characterise their genetic backgrounds. Alongside 56 publicly available genomes, we generated 479 complete ETEC assemblies from diarrhoeal isolates collected worldwide, representing the largest plasmid-resolved ETEC dataset generated to date. Using additional public data to validate our findings, we identified major gaps in current knowledge of ETEC VF distribution and plasmid diversity. Finally, comparative analysis of virulence plasmids led to the identification of eight novel putative adhesive operons, substantially expanding the known repertoire of ETEC CFs.

## RESULTS

### ETEC virulence factor determinants are predominantly plasmid-encoded

We analysed 535 complete ETEC genomes, including 479 newly generated hybrid assemblies and 56 publicly available genomes, spanning isolates collected between 1974 and 2022 across multiple continents (Extended Data Fig. 1AB, Supplementary Table 1) with strains belonging to eight distinct phylogroups (B1 (n=260, 48.6%) and A (n=210, 39.3%) being the most prevalent) (Supplementary Table 2) and 96 STs (ST443 (n=52, 9.7%), ST173 (n=34, 6.4%), ST4 (n=34, 6.4%), ST2353 (n=29, 5.4%) and ST155 (n=22, 4.1%)) (Supplementary Table 3).

The ETEC genomes displayed a diverse repertoire of CFs, toxins and non-canonical VFs. In total 21 CFs were identified (Extended Data Fig. 1C), and the most common CFs were CS6 (n=159, 29.7%), CS21 (n=151, 28.2%), and CS3 (n=68, 12.7%). The remaining CFs were present in less than 10% of all the genomes each. The most common CF profiles were CS6-only (n=71, 13.3%), CFA/I+CS21 (n=49, 9.2%), CS5+CS6 (n=47, 8.8%) and CS1+CS3+CS21 (n=33, 6.2%). Also, 68 isolates (12.7%) lacked a known CF (Supplementary Table 4).

Among the ETEC toxins linked to disease in humans, the most prevalent toxins were LT (n=362, 67.7%), STah (n=242, 45.2%) and STap (n=135, 25.2%), with LT+STah (n=125, 23.4%) and LT+STap (n=74, 13.8%) being the most common toxin combination profiles (Supplementary Table 5). Notably, STb, linked to ETEC disease in animals, was present in four genomes as the sole toxin (0.7%). Previously described non-canonical VFs EatA (n=239, 44.7%) and EtpBAC (n=207, 38.7%) were also common, present in over a third of the genomes. Shiga toxin Stx2 was present in 9 isolates (1.7%), which also harboured ETEC toxins, making them hybrid pathotypes ETEC/STEC.

The generated complete assemblies allowed us to locate the ETEC VF determinants and confirmed that all CFs, enterotoxins, and non-canonical VF genes were plasmid-encoded, apart from CS2 and *stx2* (Extended Data Fig. 1C). A total of 2,188 plasmids were identified (mean of 4.1 per isolate), of which 903 (41.3%) carried ETEC virulence factors (toxins [LT, ST], CFs, and non-canonical VFs [EatA, EtpBAC]). Plasmid size ranged from 1,000 to 263,241 base pairs (bp), with VF plasmids generally exceeding 100,000 bp, whereas non-VF plasmids were smaller (Extended Data Fig. 2A). Most plasmids were conjugative (n=850, 38.8%) or mobilisable (n=783, 35.8%), with VF plasmids more frequently classified as conjugative (Extended Data Fig. 2B).

The most common replicon type groups were IncF (n=763, 34.9%), Col-like (n=532, 24.3%), IncI (n=156, 7.1%), with most of the VF plasmids belonging to the IncF group (Extended Data Fig. 2C). Notably, 673 plasmids, including 221 carrying virulence genes, lacked an identifiable replicon, highlighting limitations of replicon-based classification^39^.

### Virulence plasmids form distinct clusters associated with colonization factors

To investigate the relatedness between plasmids we used Pling (v2.0.0)^13^. All VF plasmids belonged to a single community (Extended Data Fig. 2DEF). We then restricted the analysis to VF-positive plasmids, identifying two primary communities, C0 and C1, comprising 185 and two subcommunities, respectively. Notably, each VF plasmid subcommunity was predominantly associated with a single CF, although the same CF could be identified in multiple subcommunities (Supplementary Table 6), Extended Data Fig. 3A). To assess whether these subcommunities remain stable, we investigated the presence of the plasmids across year and country of isolation and showed that they do persist over time and are spread across multiple countries (Extended Data Fig. 4). In addition to harbouring CFs, these plasmids often harboured a toxin, but there was no evident association between toxin-profile and plasmid subcommunity (Extended Data Fig. 5).

Subcommunities associated with the same CF occupied neighbouring regions of the rearrangement-distance network (Fig. 1A). The main exception was a subset of plasmids that clustered more closely with one another despite belonging to different subcommunities. To further examine these relationships, we calculated the mean containment distance between plasmids carrying different CFs (Fig. 1B). As expected, plasmids carrying the same CF showed the lowest containment distances, as reflected by the diagonal of the matrix. Consistent with the rearrangement distance network, a subset of CFs (indicated with a star in Fig. 1B) exhibited lower pairwise containment distances to each other than the remaining CFs. Nevertheless, containment distances between plasmids carrying different CFs remained substantially greater than those observed among plasmids carrying the same CF, confirming that each CF is predominantly associated with a distinct plasmid background.

**Fig. 1:**
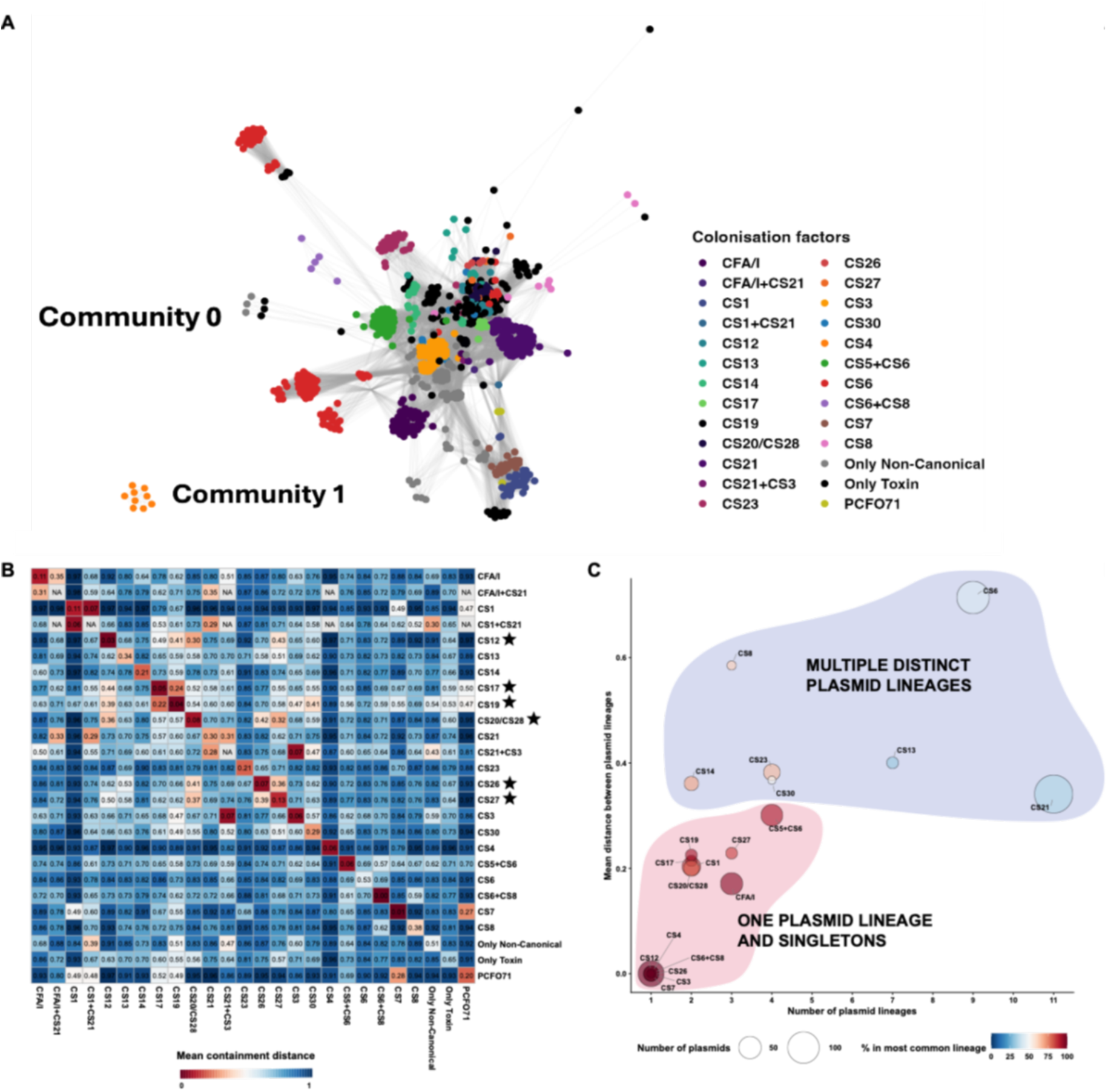
Similarity analysis of the plasmids containing virulence factors. **(A)** Rearrangement distance network of the plasmids containing VFs. The nodes are colour-coded based on the CF present in each plasmid. Plasmid communities are indicated in the figure. (**B)** Heatmap displaying the mean containment distance between the plasmids containing the different colonization factors (CFs). CF plasmids that are more similar to each other are marked with a star. (**C)** Bubble plot representing the relationship between the number of plasmid lineages a CF is present in and the mean containment distance between them. Bubble size represents the number of plasmids, and the colour indicates the percentage of plasmids belonging to the most common plasmid lineage. Highlights show the two major CF-plasmid associations. For visualisation purposes, only plasmids containing CFs present in more than 5 plasmids are shown.

Because neighbouring subcommunities carrying the same CF/CF-profile represented structurally rearranged variants of a conserved backbone, we grouped related subcommunities into higher-order plasmid lineages (see Methods). This approach grouped the 187 subcommunities into 112 plasmid lineages, including 38 singletons, with the association between CFs and plasmid groups remaining evident at the lineage level (Extended Data Fig. 3B). Notably, two main CF-plasmid associations emerged: a subset (n=12) of CFs/CF-profiles were restricted to a distinct plasmid lineage with a conserved plasmid backbone, of which six CFs were also found in plasmid lineage singletons. This is in contrast to the seven CFs which were present in different plasmid backbones across multiple distinct plasmid lineages (Fig. 1C).

### Plasmid lineages range from lineage-restricted to broadly disseminated across ETEC

Having established that CFs differ in their association with plasmid lineages, we next analysed their distribution across the ETEC core genome phylogeny (Fig. 2A). For CFs associated with only one plasmid lineage, two scenarios were observed. (i) Plasmid lineages associated with CFs CS4, CS6+CS8, CS7, CS1, and CS5+CS6 were confined to restricted regions of the core genome phylogeny, indicating clade-specific plasmids maintained predominantly by vertical descent. (ii) In contrast, CFs CS17, CS19, CS26, CFA/I, CS27, CS3, CS20/CS28, and CS12 were detected across distant branches of the ETEC phylogeny.

**Fig. 2:**
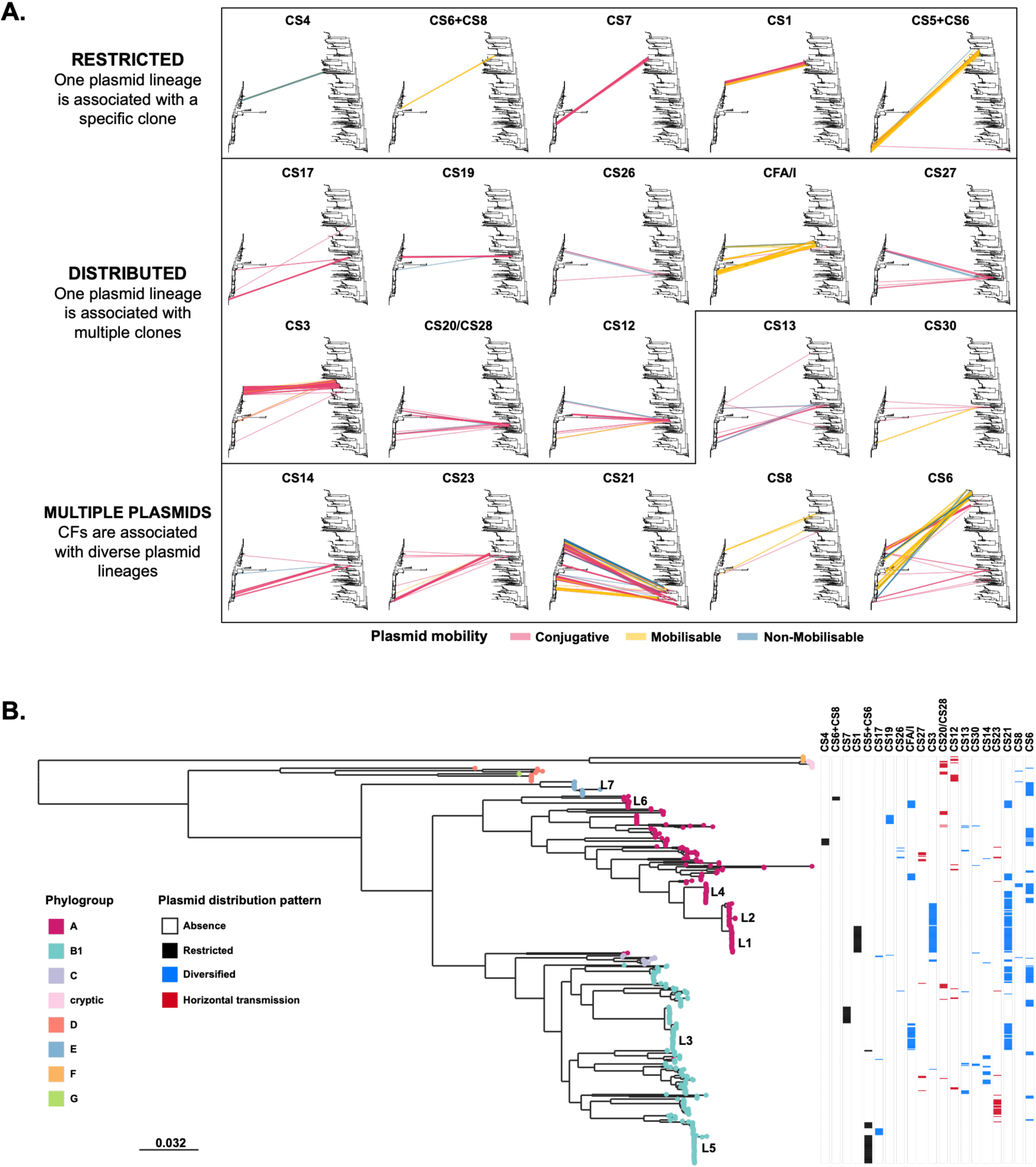
Plasmid distribution across the ETEC core phylogeny. **(A)** Tanglegrams showing the distribution of virulence factor (VF) plasmids across the core genome phylogeny. Trees to the left represent an ETEC maximum-likelihood core genome phylogeny and the tree to the right is a neighbour-joining plasmid phylogeny based on the pling containment distances. The connecting lines are coloured based on predicted plasmid mobility. CFs are ordered according to the maximum core genome distance between the distinct positive isolates to each CF. Only CF combinations with more than 5 isolates are shown. (**B)** An ETEC core genome phylogenetic tree showing the presence/absence of each combination of CF. Block colours indicate plasmid distribution patterns: black, restricted; blue, diversified; and red, patterns suggesting horizontal transmission.

A subset of CFs was associated with multiple distinct plasmid lineages, but the plasmids themselves were phylogenetically restricted (CS13, CS30). In contrast, other CFs (e.g. CS14 and CS23) were carried by multiple plasmid lineages broadly distributed across the phylogeny. Another group (CS21, CS6, and CS8) displayed mixed patterns, containing plasmid lineages that were restricted and others distributed (maximum phylogenetic breadth larger than 0.1) (Extended Data Fig. 6A).

To investigate whether these broadly distributed plasmid lineages reflected dissemination of individual plasmids or multiple phylogenetically restricted plasmid variants, we analysed these patterns at the Pling subcommunity level (Extended Data 6B) and showed that most of the subcommunities were phylogenetically restricted, suggesting that these lineages could have reached a similar genetic content through evolution patterns and then be vertically transmitted.However, plasmids carrying CS12, CS20/CS28, CS23, and CS27 (each associated with conjugative plasmids) were distributed across distant phylogenetic lineages, providing strong evidence of horizontal transfer in these cases.

Taken together, analyses revealed three evolutionary models underlying the distribution of ETEC CFs (Fig. 2B): (i) phylogenetically restricted CFs maintained within specific ETEC lineages, (ii) a suggested convergent or long-term coevolution of the same CF present in distinct plasmid subcommunities associated with a specific ETEC clade, and (iii) broadly distributed CFs present on conjugative plasmids suggesting horizontal dissemination.

### Novel adherence factor operons expand the repertoire of ETEC colonization factors

Because a substantial proportion of toxin-positive isolates lacked recognized CFs (12.7%, 68/535) (Supplementary Table 4), we next asked whether additional adhesins followed the same evolutionary framework established above. Furthermore, because the most of CF-carrying plasmids also contain ETEC toxins (Extended Data Fig. 5), plasmids carrying only toxins might harbour novel adherence factors. Therefore, we explored Bakta^14^ annotations, and InterPro^15^ functional predictions, to identify classical chaperone-usher-fimbrial subunit CF operon structures.

We identified eight novel putative adherence factor operons (Supplementary Material) which belonged to the γ-, κ-, and π-fimbrial usher protein (FUP) clades (Fig. 3A). The operons were provisionally named according to their phylogenetic placement.

**Fig. 3.**
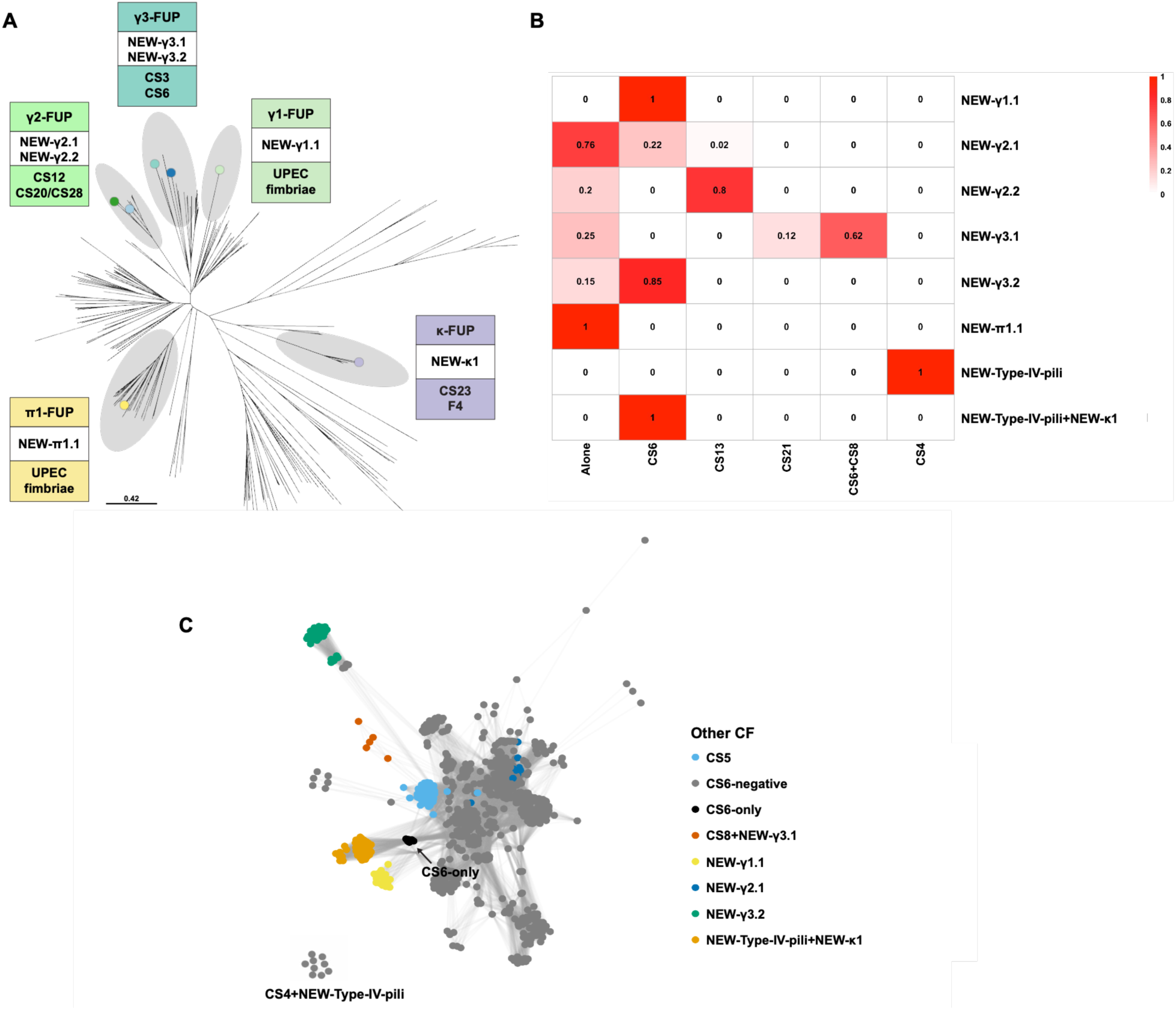
Phylogenetic and plasmid context of novel putative ETEC adherence factors. **(A)** Phylogenetic tree based on an alignment of 189 previously described fimbrial usher protein sequences from Nuccio et al^16^. Coloured nodes indicate usher sequences identified in the novel putative CF operons described in this study. FUP clades containing these sequences are circled. Boxes indicate the FUP clade, the corresponding putative CF operons identified here, and examples of previously characterised CFs belonging to the same clade. (**B)** Heatmap displaying the proportion of other known CFs on the same plasmid as a new putative adherence factor. Values closer to 0 (absence) are shown in white and closer to 1 (presence in all plasmids) are shown in red. (**C)** rearrangement distance network of the plasmids containing VFs with nodes coloured with the putative/known CFs coexisting in the same plasmid. The only plasmid cluster that did not contain any other CF is labelled in the figure. However, these isolates harboured another plasmid containing CS4 and the NEW-Type-IV-pili.

Given the strong associations observed between known ETEC CFs, plasmid lineages and ETEC phylogeny, we next examined whether the predicted adherence factor operons followed similar patterns. Like recognized CFs, the putative operons were associated with distinct plasmid subcommunities and plasmid lineages and showed structured distribution across the ETEC core phylogeny (Extended Data Fig. 7). To further assess their potential role in ETEC pathogenicity, we examined their prevalence in the AllTheBacteria (ATB) database. Most of these CFs were not rare, except for NEW-γ3.1, and they were predominantly associated with ETEC toxin-positive isolates, supporting their role in ETEC pathogenicity (Extended Data Fig. 8).

Notably, these putative adherence factors were not always present as the sole adhesive determinant, but frequently co-occurred with established CFs in the same isolate (Fig. 3B). This was particularly evident for CS6, one of the most prevalent CFs. Most of the plasmid lineages carrying CS6 also encoded either a known CF (CS5) or a new putative adherence factor (NEW-γ3.1+CS8, NEW-γ1.1, NEW-γ2.1, NEW-γ3.2 or NEW-Type-IV-pili+NEW-κ1). Only a single plasmid cluster (labelled in Fig. 3C) contained CS6 on its own, but the same isolates contained CS4+NEW-TypeIV-pili in another plasmid, suggesting that CS6 tends to be accompanied by another CF (Fig. 3C).

### Plasmid lineage patterns are largely conserved but reveal additional diversity in public data

The CF-associated plasmid lineage-host associations identified in the complete ETEC genome collection were largely recapitulated in the broader *E. coli* context (Fig. 4A). For most CFs, the same plasmid lineage remained associated with the same dominant STs, indicating that the major plasmid–host associations observed in our dataset were robust. The larger ATB dataset also clarified several cases that appeared ambiguous in the complete ETEC genome collection. For example, CS5+CS6 and CS17 were associated with individual plasmid lineage singletons in our dataset, and screening ATB confirmed that these plasmid lineages remained strongly associated with the same dominant STs. Similarly, the CS19 plasmid lineage associated with ST3864, represented by a single isolate in the complete ETEC genome collection, was repeatedly identified in ATB, demonstrating that this lineage was not an isolated event.

**Fig. 4:**
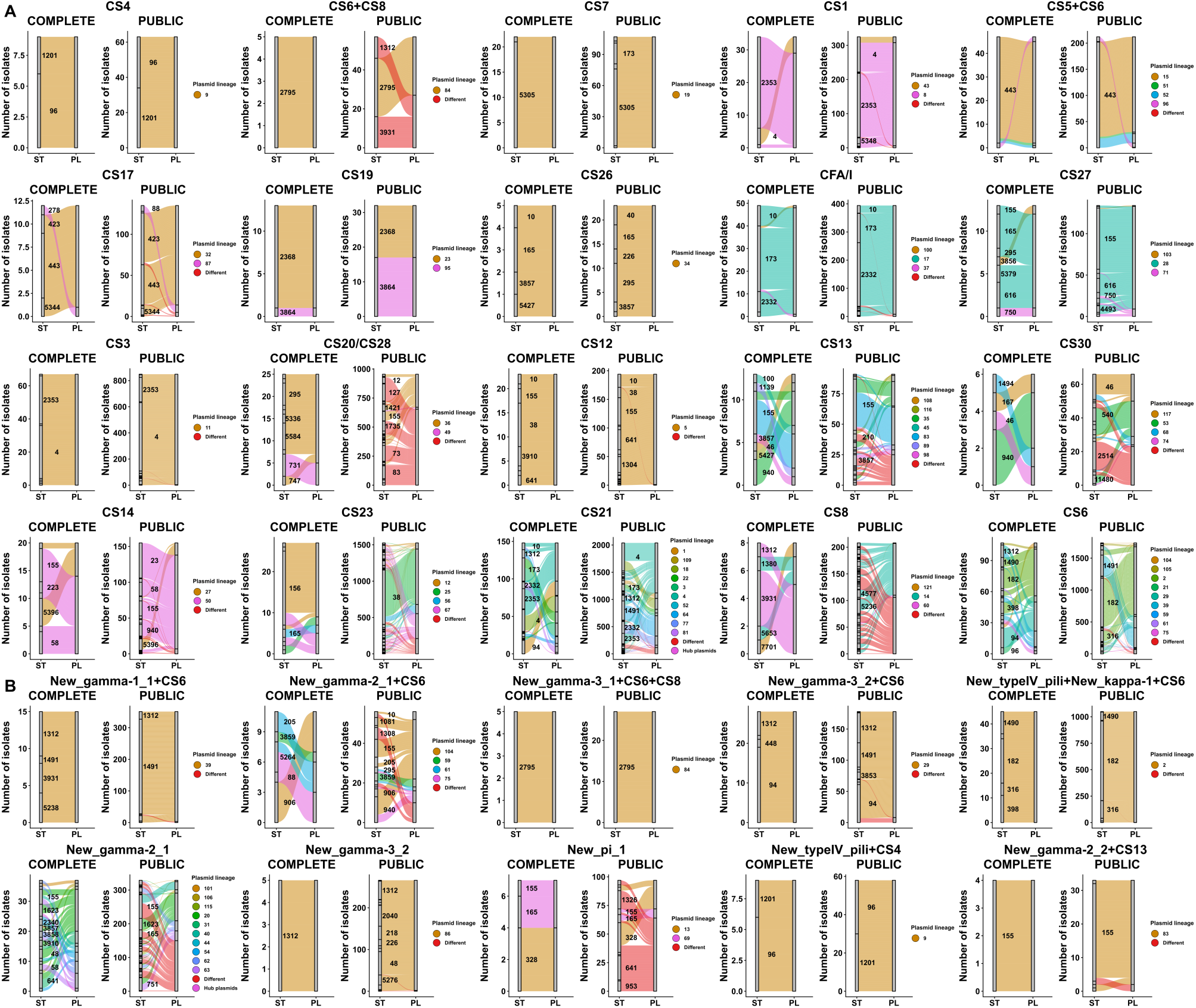
Validation of the plasmid distribution findings using ATB *E. coli* genomes. Alluvial plots displaying the distribution of each plasmid lineage into the different STs in complete assemblies generated in this study and short-read assemblies publicly available in ATB dataset for the known CFs (**A)** and the putative newly discovered adhesive operons (**B**) present in more than 5 isolates in the dataset generated in this study. Left stratum represents Sequence Type (ST), and the right stratum represents Plasmid Lineage (PL). Each colour represents a distinct plasmid lineage; red indicates that no defined plasmid from this study was detected. For visualisation purposes, only STs present in more than 5% of the genomes containing a specific CF were labelled.

A subset of CF-associated plasmid lineages differed from patterns observed in the complete ETEC genome dataset. CS20/CS28 and CS23 isolates were mostly toxin-negative and associated with STs in which enterotoxin-positive isolates are absent (Extended Data Fig. 9). One exception was ST38 carrying CS23, linked to plasmid lineage 25 that lacked toxin genes, raising the possibility of a transition from non-ETEC to ETEC backgrounds. In contrast, CS13 and CS30 were mostly found in ETEC isolates, pointing instead to missing plasmid diversity in the complete ETEC genome dataset. CS8 showed a more mixed pattern, with a group of unknown plasmids associated with ETEC isolates and another with non-ETEC.

To determine whether the putative adherence factor operons followed similar evolutionary patterns to known ETEC CFs, we analysed their distribution in the ATB dataset (Fig. 4B). Like established ETEC CFs, the putative adherence factor operons displayed both lineage-restricted and broadly distributed plasmid associations. The ATB dataset further showed that some putative operon-associated plasmid-lineages were present in additional *E. coli* STs not represented in our complete genome dataset (e.g. NEW-γ2.1+CS6 and NEW-γ3.2). Furthermore, many isolates with putative adherence factor operons lacked detectable known plasmids (e.g. NEW-γ2.1 and NEW-π1), making them attractive candidates for future long-read sequencing if confirmed as functional adhesins.

### CFs linked to horizontally transmitted plasmids have disseminated beyond classical ETEC clonal backgrounds

To investigate whether dissemination of individual CFs was associated with particular *E. coli* populations, we examined the distribution of CF-positive genomes across sequence types in the ATB dataset.

To distinguish CFs largely confined to the classical ETEC population from those occurring more broadly, we classified sequence types according to the proportion of toxin-positive genomes in the ATB dataset (see Methods). For each CF, we quantified its prevalence across STs together with the proportion of toxin-positive genomes within each ST, allowing the distribution of individual CFs to be interpreted in the context of the broader *E. coli* population.

Those CFs described as major, such as CFA/I, CS3, and CS6^6^, remained predominantly confined to ETEC-associated STs, even when detected in multiple STs (Fig. 5). In contrast, CFs associated with horizontally transmitted plasmids (Fig. 2B) (including CS23, CS20/CS28, CS12 and CS27), as well as CS13 and CS30, which presented a broader phylogenetic pattern in the ATB dataset (Fig. 4), were mostly detected in non-ETEC specialized STs. These findings indicate that a subset of ETEC CF-associated plasmids circulate in a broader range of *E. coli* genetic backgrounds than the classical ETEC-associated CFs.

**Fig. 5:**
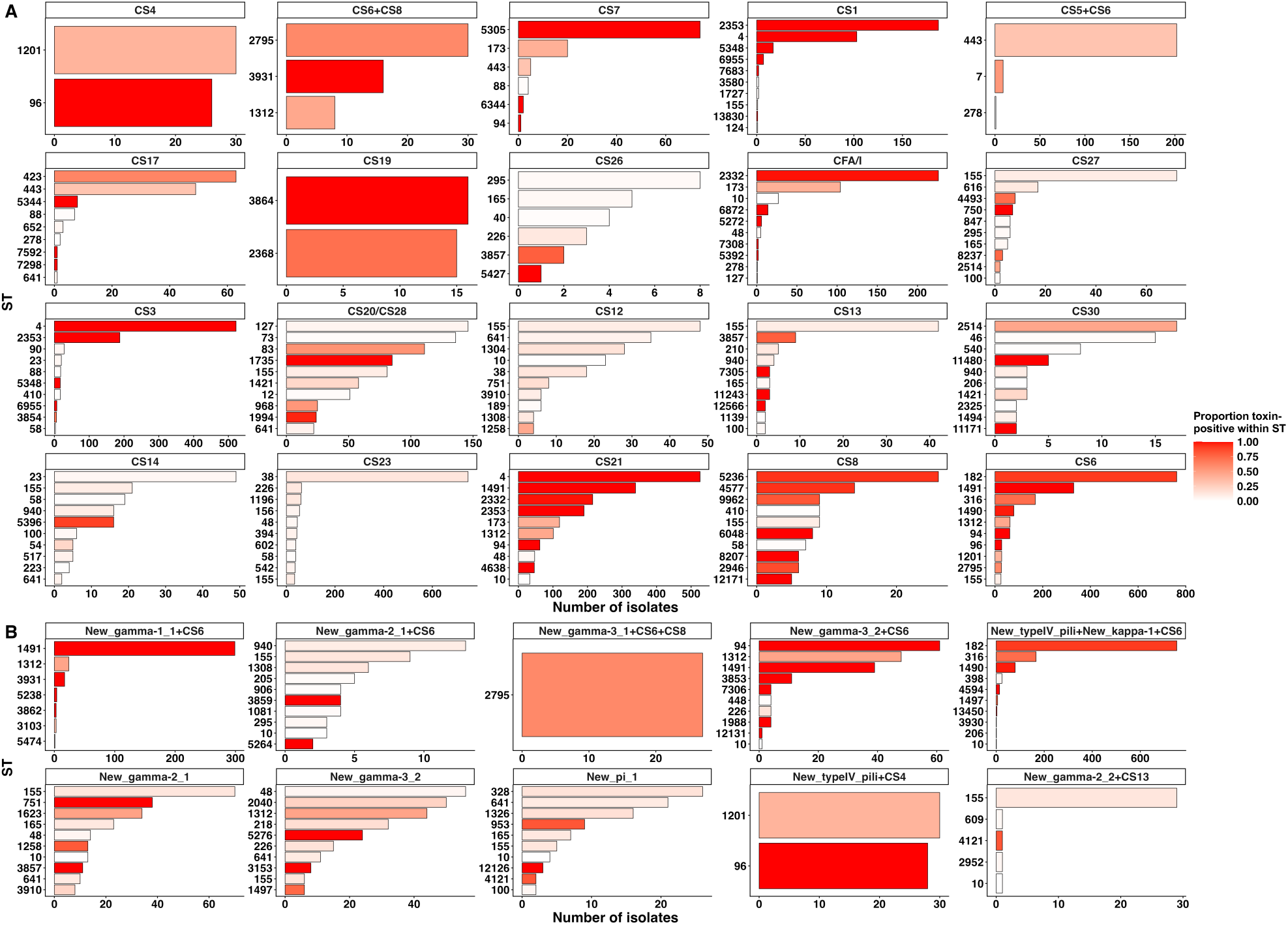
CF association with ETEC or non-ETEC clones. Distribution of CFs across sequence types with differing proportions of toxin-positive isolates. Bar plots show the up to 10 most frequently represented sequence types (STs) among ATB isolates carrying **(A)** known CFs or CF combinations and **(B)** putative adherence operons or operon combinations identified in more than five isolates in our study collection. Bar length indicates the number of isolates carrying the indicated profile within each ST. Bar colour indicates the proportion of all isolates belonging to that ST in ATB that carry at least one ETEC enterotoxin gene, ranging from white (0) to red (1). This proportion was calculated using all isolates of each ST, irrespective of their CF profile.

The CF CS6, apart from being the most prevalent CF in most epidemiologic studies, is especially interesting since each plasmid was associated with another known or new predicted adherence factor (Fig. 3C). Even though it is mainly located in ETEC specialist STs, a subset of them were detected outside classical ETEC genetic backgrounds (Fig. 5A). Interestingly, we showed that most of them were isolates that also contained also the NEW-γ2.1 (Fig. 5B), suggesting a possible impact for its epidemiology. Finally, when the new putative adherence operons were found alone, they were detected in non-ETEC STs, indicating that they would behave more similarly to the non-classical than to the classical CFs.

## DISCUSSION

In this study, we generated the largest human ETEC complete genome collection to date, with 479 newly generated assemblies, representing a substantial increase compared with the 56 complete genomes available at the time of this study. By analysing this collection, we described distinct plasmid distribution patterns associated with ETEC cases, validated these findings using the vast amount of available short-read data and identified priorities for future ETEC research and surveillance.

The key role of plasmids in ETEC virulence was confirmed, with three major distribution patterns being identified: (i) one plasmid lineage restricted to a single ETEC clade, (ii) one plasmid lineage distributed across multiple ETEC clades and (iii) multiple plasmid lineages distributed across multiple phylogenetic clades. We showed that the plasmid lineages with the broadest distribution tended to be predicted as conjugative. This partly supports the finding that major ETEC lineages drive most clinical cases, as reported in the foundational ETEC genomics study^9^. However, for many of the most prevalent CFs, our results indicate that plasmids, rather than bacterial lineages, represent the most informative unit of study to understand ETEC dissemination, as has already been proposed for AMR dissemination^17^.

The Pling clustering tool^13^ represents a major advance in the field because it not only classifies plasmids according to their genomic context but also incorporates structural events, i.e. pling’s rearrangement distances. For CS23, CS12 and CS20/CS28, plasmids belonging to the same Pling subcommunity occurred in phylogenetically divergent ETEC hosts (Extended Data Fig. 6B). The occurrence of closely related plasmids across these host backgrounds suggests horizontal dissemination. In contrast, other major CFs, such as CS3 and CFA/I, were associated with only one plasmid lineage but displayed substantial structural differences while retaining a conserved plasmid backbone. This suggests that these plasmids have diversified extensively following acquisition rather than representing recent horizontal transfer. Whether the diversification reflects long-term maintenance, plasmid remodelling or convergent evolution remains to be determined. These findings suggest that certain CFs may require specific plasmid backgrounds to be maintained successfully and contribute to pathogenesis, even if they are not the same plasmid. This observation is particularly relevant for ETEC control strategies, as it raises the possibility of alternative interventions based on plasmid curing^18^, which could have a lower impact on the microbiome and bacterial ecology than conventional antimicrobial approaches, which could also be applied in environmental settings.

Taking advantage of the strong association between plasmid lineages and host STs, we validated these patterns by mapping plasmid lineages to publicly available short-read assemblies. This analysis confirmed most of the patterns observed in the complete genome collection, while also highlighting important gaps in current plasmid knowledge, suggesting that additional CS20/CS28-, CS13- and CS30-positive isolates should be sequenced to complete the picture of plasmid diversity in ETEC. Furthermore, placing these findings within the broader *E. coli* context revealed two distinct ecological patterns. Classical major CFs, such as CS6, CFA/I, and CS3, were predominantly associated with specialist ETEC clones, whereas other less prevalent CFs^6,9^, including CS12, CS20/CS28 and CS23, were found beyond the classical ETEC genetic backgrounds and were common in the ATB dataset. Unfortunately, metadata were not available for all isolates, making it impossible to determine whether this pattern has existed historically or emerged more recently. Notably, these CFs were associated with conjugative plasmids whose distribution across distant ETEC lineages is consistent with horizontal dissemination, suggesting that conjugation has facilitated their spread into non-classical genetic backgrounds.

Although the biological basis of this pattern remains unclear, it opens a new line of research into the ecology and evolution of ETEC. Classical CFs have historically been associated with the most severe cases of disease^6^, but these studies were largely based on PCR assays in which many of the less common or recently described CFs were not included. However, CS23, a CF that has recently gained interest due to its increasing in Bangladesh, has been shown to be associated with severe disease^10^, suggesting that CFs currently considered minor may contribute substantially to disease burden. The presence of these CFs outside of classical ETEC STs raises important questions regarding their pathogenic potential and highlights the need for studies that integrate CF identity, plasmid background and chromosomal lineage when assessing disease severity^19^. These findings also raise questions regarding transmission routes. One possibility is that classical CFs, present in ETEC lineages, are primarily maintained through human-to-human transmission, whereas broader phylogenetic distribution of CFs on conjugative plasmids suggests that their ecological niche may extend beyond ETEC populations. Unfortunately, environmental studies investigating CF distribution remain uncommon. To our knowledge, only one such study has been performed in Bolivia^20^, where CS23 was the most prevalent CF identified in water samples and was associated with ST410, a non-ETEC specialist lineage. Further investigation of environmental reservoirs will therefore be required to better understand the transmission cycle of ETEC.

Important gaps also remain in ETEC epidemiology, as a substantial proportion of toxin-positive isolates do not carry any known CF^5^. Considerable efforts have been made to identify new CFs, and several have been described in recent years^21^. Most of these discoveries have relied on bioinformatic approaches based on sequence similarity to known CFs^22^. Although this strategy has successfully identified new putative CFs, many have not yet been experimentally validated^23,24^, while for others the gene sequences have not been publicly released^25^. Taking advantage of complete genome assemblies, we observed that plasmids carrying known CFs were almost invariably associated with enterotoxin genes. We therefore reasoned that toxin-positive plasmids lacking recognized CFs represented promising targets for the identification of novel adhesins. Through InterPro^15^ annotation of these plasmids, we identified additional chaperone-usher families not represented among currently recognized ETEC CFs, including members of the π- and γ1-FUP clades. However, when analysed in the context of the global *E. coli* population, these putative CFs were predominantly associated with toxin-positive isolates (Extended Data 7), and those that occurred without any additional CF displayed distribution patterns similar to those observed for CFs present in non-ETEC *E. coli* (Fig 5B). Experimental validation will be required to confirm their role in colonization. Together, these observations support the hypothesis that these loci could represent genuine CFs and suggest that their validation could substantially reduce the current gaps in ETEC epidemiology.

One particularly interesting finding involved CS6, one of the most common CFs globally^6^. Its association with CS5 is already well established^26^. However, when putative CFs were also considered, every CS6-positive isolate contained at least one additional CF, and each combination was associated with a distinct plasmid lineage. These observations are especially relevant because CS6 is one of the main targets of several ETEC vaccine candidates currently under development^27^. The presence of additional adhesins may influence immune recognition and vaccine effectiveness. Consequently, the biological and immunological consequences of this diversity should be investigated in greater detail.

Our results show that plasmids are a key determinant of how ETEC CFs are maintained and spread, confirming stable lineage-restricted associations while revealing broader plasmid-driven dissemination of specific CFs across diverse *E. coli* backgrounds. Plasmid background should therefore be considered alongside clonal lineage in ETEC surveillance. We additionally identified eight novel putative adherence factor operons that may explain why many toxin-positive ETEC isolates lack recognized CFs, expanding the candidate repertoire of ETEC CFs.

## MATERIAL AND METHODS

### Bacterial isolates, DNA extraction and genome sequencing

A total of 479 ETEC isolates from the Svennerholm and von Mentzer UGOT ETEC collection were selected for sequencing. The isolates were obtained from diarrhoeal cases in low- and middle-income countries between 1980 and 2014. To complement the newly generated genomes, publicly available ETEC genomes were identified through NCBI Pathogen Detection. Genomes with fewer than 10 contigs and available metadata regarding source, country of origin, and year of isolation were screened for the presence of enterotoxin genes (LT, STah, STap, or STb). Fifty-six genomes fulfilled these criteria and were included, resulting in a final dataset of 535 ETEC genomes.

For newly sequenced isolates, strains were cultured overnight on horse blood agar plates at 37°C to verify purity. Genomic DNA for short-read sequencing was extracted using the Wizard Genomic DNA Purification Kit (Promega), while DNA for long-read sequencing was extracted using the FireMonkey High Molecular Weight DNA kit (Revolugen), according to the manufacturer’s instructions. The DNA concentration was measure using a NanoDrop spectrophotometer and the DNA quality was analysed using the 4150 TapeStation System and samples with DIN > 9 were selected for sequencing.

Short-read sequencing was performed using Illumina platforms. A subset of isolates had previously been sequenced on the Illumina HiSeq 2000 platform with 75 or 100 bp paired-end reads, while the remaining isolates were sequenced on the Illumina NovaSeq platform with 150 bp paired-end reads. Libraries were prepared according to this protocol^28^, and unique index sequences were used for multiplexing.

For the majority of isolates, long-read sequencing libraries were prepared from up to 400 ng DNA using Oxford Nanopore Technologies ligation sequencing kit (SQK-LSK109) with native barcoding (EXP-NBD196). Library preparation included end repair, barcode ligation, SPRI bead purification, pooling and adapter ligation. Libraries were sequenced on PromethION flowcells, and reads were basecalled using Guppy^29^ (v6.0.6).

For the remaining isolates, long-read sequencing libraries were prepared using Rapid Barcoding Kits (SQK-RBK004 or SQK-RBK114-24), with up to 200 ng DNA per sample and three samples multiplexed per run. Libraries were sequenced on a MinION Mk1B using R10.4.1 Flongle flow cells (Oxford Nanopore Technologies). Real-time basecalling used the high-accuracy model, and reads were filtered using a minimum length of 1,000 bp and a minimum Q score of 9.

### Hybrid assemblies

Long reads generated were combined with the short reads using Unicycler (v0.5.1, default parameters)^30^. Quality metrics of the generated assemblies were determined using QUAST (v5.3.0)^31^ and are available in Supplementary Table 1. To confirm that the resulting assemblies were suitable for the intended plasmid analysis, the presence of ETEC VFs was screened against a customized dataset (https://github.com/avonm/ETEC_vir_db)^32^ with Abricate^33^ (v1.4.0, default parameters) and only those isolates with positive hits in a closed contig were selected.

### Core-genome phylogenetic analysis and bacterial annotation

*E. coli* hybrid assemblies were annotated using Bakta (v1.12)^14^. Core and accessory genome were determined using Panaroo (v1.17)^34^, with the core genome defined as genes present in at least 95% of isolates. Core gene alignments were generated using MAFFT (v7.526)^35^. The concatenated core gene alignment was used as an input for IQ-TREE^36^ (v3.0.1, default parameters, using substitution model GTR-GAMMA) to build a maximum-likelihood core-genome phylogenetic tree. The tree was visualized and annotated with Microreact^37^ and can be accessed here: https://microreact.org/project/sgbYhmtLLYL9f2Yg11yth2-ecolietecallvf.In parallel, genomes were assigned to phylogroups using ClermonTyping (v23.06)^38^ and grouped into sequence types (ST) with MLST (v2.35.0)^39^ according to the seven gene Achtman scheme^40^.Abricate^33^ (v1.4.0) was used to scan genomes against a customized dataset (https://github.com/avonm/ETEC_vir_db)^32^ to identify CFs, enterotoxins, Shiga-toxins and non-canonical ETEC VF encoding genes. A positive hit was considered with a minimum identity and minimum coverage of 80%. This information together with the metadata available is summarized in Supplementary Table 1.

### Plasmid analysis

Near-complete assemblies allowed us to consider a contig as plasmid or chromosomal origin according to its size, but in order to confirm it, we used plasmidEC (v1.3.1)^41^, a binary plasmid classification tool which combines Centrifuge^42^, Platon^43^ and RFPlasmid^44^, resulting in a total of 2,188 plasmids. All these contigs were again scanned with Abricate^33^ (v1.4.0) against the customized ETEC dataset as previously described. When a positive hit was detected, it was considered an ETEC virulence plasmid, resulting in 903 ETEC virulence plasmids. Mobility status was determined with the MOB-Suite (v3.1.9)^45^ software with the MOB-Typer option. Replicon types were determined with Abricate, scanning the plasmids against PlasmidFinder^46^ dataset. All this information is summarized in Supplementary Table 7.

### Plasmid clustering and lineage definition

All plasmid contigs were clustered with Pling (v2.0.0)^13^. Pling combines containment distances and double cut and join (DCJ)-Indel rearrangement distances to identify plasmid communities and subcommunities representing broad and close plasmid relatedness, respectively^13^.

As the primary focus of this study was the distribution of ETEC VFs, subsequent analyses were restricted to plasmids carrying ETEC VFs. Initial clustering was performed using the default Pling settings (containment distance = 0.5 and double cut and join distance threshold = 4), resulting in plasmid communities and subcommunities. Summary information about VF plasmids can be found in Supplementary Table 8.

Because the dataset included isolates collected over multiple decades and continents, Pling subcommunities were highly granular. To capture broader evolutionary relationships among virulence plasmids, higher-order plasmid lineages were generated by Pling by hierarchical clustering of the pairwise containment distances. Subcommunities with a containment distance of ≤ 0.15 were grouped into the same plasmid lineage. The resulting plasmid lineages are summarized in Supplementary Table 7.

The clustering was selected by comparing representative plasmids generated across a range of containment distance thresholds (0.1, 0.15, 0.2 and 0.25). Representative CF-associated plasmids of each subcommunity were visualized using Clinker^47^, and the threshold showing the best agreement between hierarchical clustering and conserved plasmid backbone was selected. Representative comparisons are provided in the Supplementary Material.

For visualisation, a neighbour-joining plasmid tree was constructed from the pairwise containment distance matrix using rapidnj (v2.3.3)^48^. Phylogenetic breadth of each plasmid lineage and subcommunity were calculated with treedist^49^ and those with a maximum breadth greater than 0.1 were considered distributed.

### Identification of new putative adherence factors

All annotated ETEC VF plasmids were screened for gene clusters encoding potential adherence factors, searching for the presence of the minimal CF gene cluster structure, i.e., clusters with at least a chaperone, an usher and a fimbrial subunit based on the InterPro protein function prediction in the Bakta annotation. The only operon outside of this structure was a type IV pilus operon with a subset of genes similar to CS8. For every individual gene, the amino acid sequences from all isolates were extracted, and the genes were clustered by similarity. Briefly, the genes with the same label annotation in Bakta were extracted and those with similarities higher than 90% were considered the same gene.

Each predicted adherence factor was assigned to a fimbrial usher protein (FUP)-clade by extracting the amino acid sequence of each putative usher and concatenating and aligning them using MAFFT (v7.526)^35^ together with a reference dataset of 189 usher proteins described by Nuccio et al^16^ and all described CFs (https://github.com/avonm/ETEC_vir_db). Next, a maximum likelihood phylogenetic tree was built with IQ-TREE (v3.0.1, default parameters, using substitution model WAG+F+G4)^36^. As the putative CFs described in this study have not been experimentally validated, the existing ETEC CF nomenclature was not followed. Instead, predicted adherence factors were named according to the usher family they belonged to. The nucleotide sequence of each gene for each putative adhesive molecule can be found in Supplementary Material and in the following GitHub repository: https://github.com/vonMentzerLab/ETEC_plasmidome.

### Retrieval of short-read ETEC public genomes and plasmid prediction

To validate our findings in a broader genomic context, all assemblies available in the ATB^50^ database (May 2025 release with 2,763,297 assemblies) were screened for the presence of the distinct CFs using LexicMap (v0.9.0)^51^ with a customized dataset as reference (https://github.com/avonm/ETEC_vir_db)^32^. Assemblies were subjected to quality control, retaining only genomes with sizes between 4 and 6.5 million bp, less than 500 contigs, and an N90 greater than 10,000 bp, resulting in 7,231 *E. coli* genomes containing a CF.

To predict the presence of plasmid lineages identified in this study, genomes were grouped according to the CF detected. For each CF-specific group, all plasmids carrying that CF in the complete genome dataset were used as references and compared against the short-read assemblies using Quast (v5.3.0)^31^. The reference plasmid with the highest genome fraction was considered the most likely plasmid carrying the CF and was assigned to a plasmid lineage according to the Pling classification described above.

Given the contig fragmentation inherent to short-read assemblies, a relaxed genome fraction threshold of 75% was applied. Assemblies with lower values were interpreted as carrying either a distinct plasmid not represented in the complete genome dataset or a chromosomally encoded CF.

All the *E. coli* genomes present in ATB meeting the quality control parameters used for this study (sizes between 4 and 6.5 million bp, less than 500 contigs, and an N90 greater than 10,000 bp) were scanned using MLST^39^ according to the seven gene Achtman scheme^40^ to determine the ST. Additionally, all the resulting genomes were screened using LexicMap (v0.9.0)^51^ with a customized dataset as reference (https://github.com/avonm/ETEC_vir_db)^32^ to identify the presence of enterotoxins. Finally, ETEC ratio within each ST was calculated as the proportion of isolates carrying an enterotoxin among all isolates assigned to that ST.

## Data availability

The sequencing data and genome assemblies generated in this study have been deposited in the NCBI repositories under BioProject accession PRJEB127140. Previously generated Illumina sequencing reads are available under their respective accession numbers, as detailed in Extended Data. Newly generated Oxford Nanopore sequencing reads and hybrid genome assemblies are currently held under private access and will be made publicly available upon publication. Access to the deposited data for peer review can be provided upon request.

## Code availability

All the scripts needed to reproduce the figures are available at the following Github repository: https://github.com/vonMentzerLab/ETEC_plasmidome.

## Acknowledgements

We thank the many researchers, clinical and laboratory teams, and collaborating institutions whose contributions since the 1980s have enabled the establishment, maintenance and continued development of the UGOT ETEC Strain Collection. We are particularly grateful to Professor Ann-Mari Svennerholm for her longstanding leadership in building and sustaining this resource.

## Funding

Astrid von Mentzer was supported by the SciLifeLab & Wallenberg Data Driven Life Science (DDLS) Program (KAW 2020.0239), the Swedish Research Council (Vetenskapsrådet; 2022-01449), the Jeanssons Foundation (J2023-0095) and Åke Wibergs Stiftelse (M22-0180). Sequencing performed at the Quadram Institute was supported by institute core funding.

## Contributions

A.v.M. conceived and supervised the study, secured funding, and contributed to data analysis and manuscript writing. B.G. performed the analyses, prepared the figures and drafted the manuscript. J.V. and R.E. performed DNA extraction. A.P. secrued funding from Quadram Institute for sequencing and contributed to manuscript writing. Z.I. and D.F. provided expertise on Pling and contributed to discussions of the plasmid analyses and manuscript review. S.D. contributed in genome assembly analysis and manuscript writing. M.G. contributed to the databases used for virulence profiling and to manuscript writing. T.L.V. and D.B performed sequencing. E.J. contributed to sequencing and manuscript writing.

## Notes

### Competing Interest Statement

The authors have declared no competing interest.

