## Extended data for "Contrasting plasmid–host associations shape the distribution of enterotoxigenic *E. coli* (ETEC) colonization factors"

### Extended Data Figures

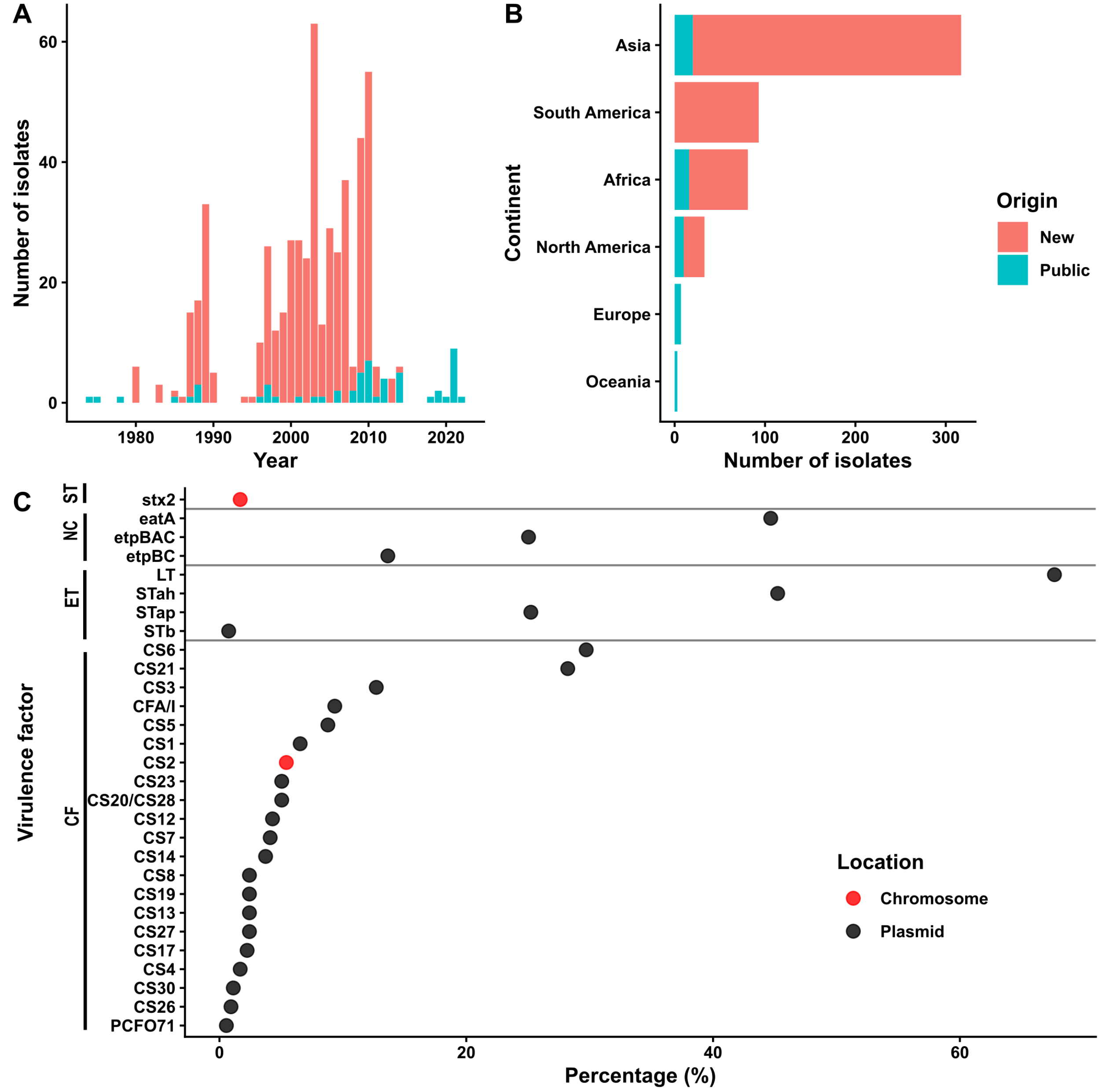


**Extended Data Fig. 1.** (A,B):Bar chart showing the number of isolates per year (A) and continent (B) coloured based on whether the complete genome was newly generated (red) or from public datasets (blue). (C): Dot plot showing the percentage of virulence factors in ETEC isolates included in the study, grouped by class (CF: colonisation factors, ET: enterotoxin, NC: non-canonical, ST: Shiga-toxin.) Points are coloured by location, either in plasmids (black) or chromosome (red).

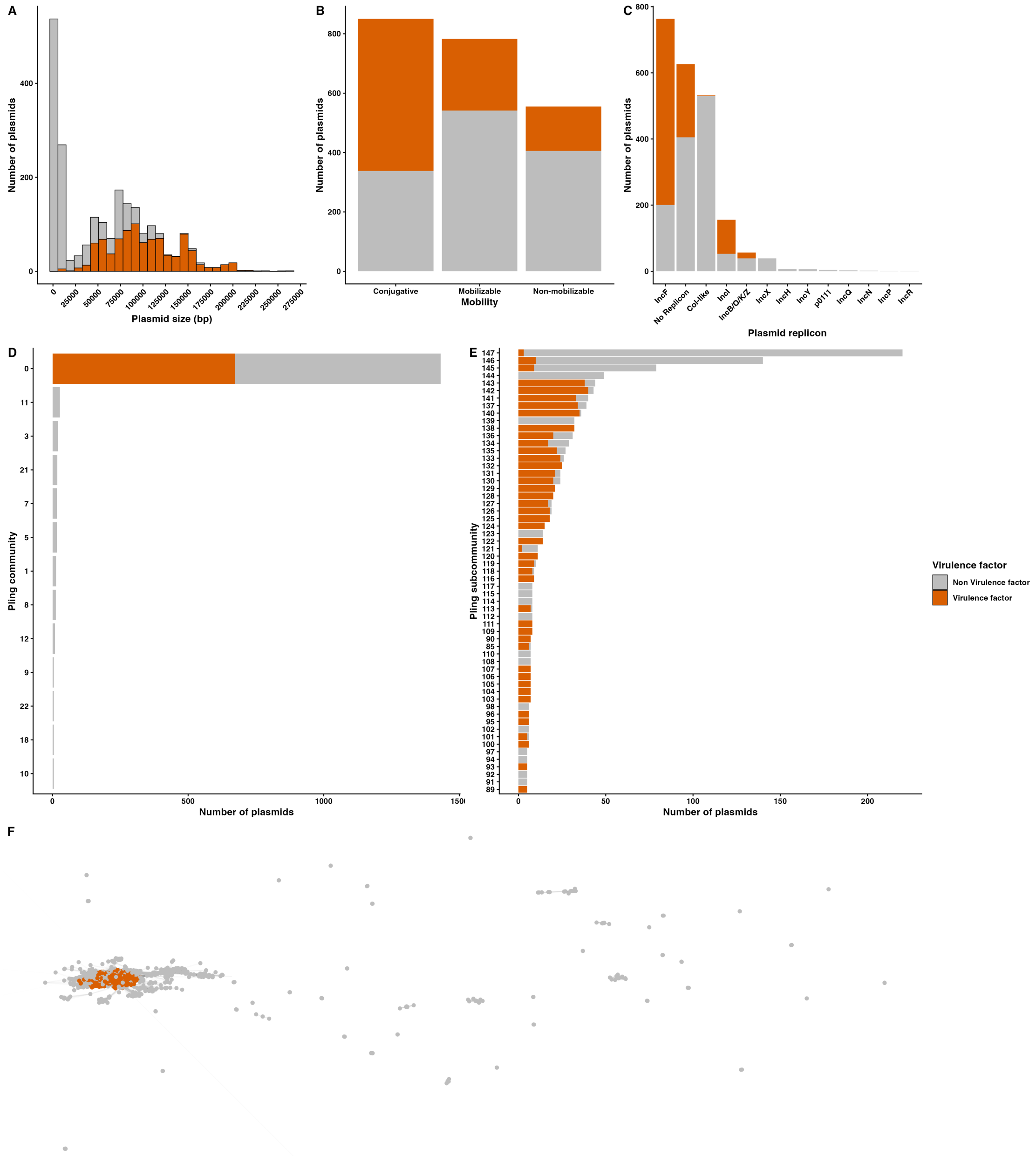


**Extended Data Fig. 2: General information about all the plasmids obtained in the study.** Grey: Plasmid that does not contain a virulence factor, orange: plasmid that contains a virulence factor. (A): Histogram showing the length of the plasmid. Bp: base pairs. (B, C, D, E): Bar charts showing the number of plasmids predicted as conjugative, mobilizable or non-mobilizable (B), the frequency of distinct plasmids replicon type groups (C), the frequency of distinct plasmid communities according Pling (D) and the subcommunities belonging to community 0 (E). (F): Rearrangement distance network of all the plasmids present in the ETEC assemblies.

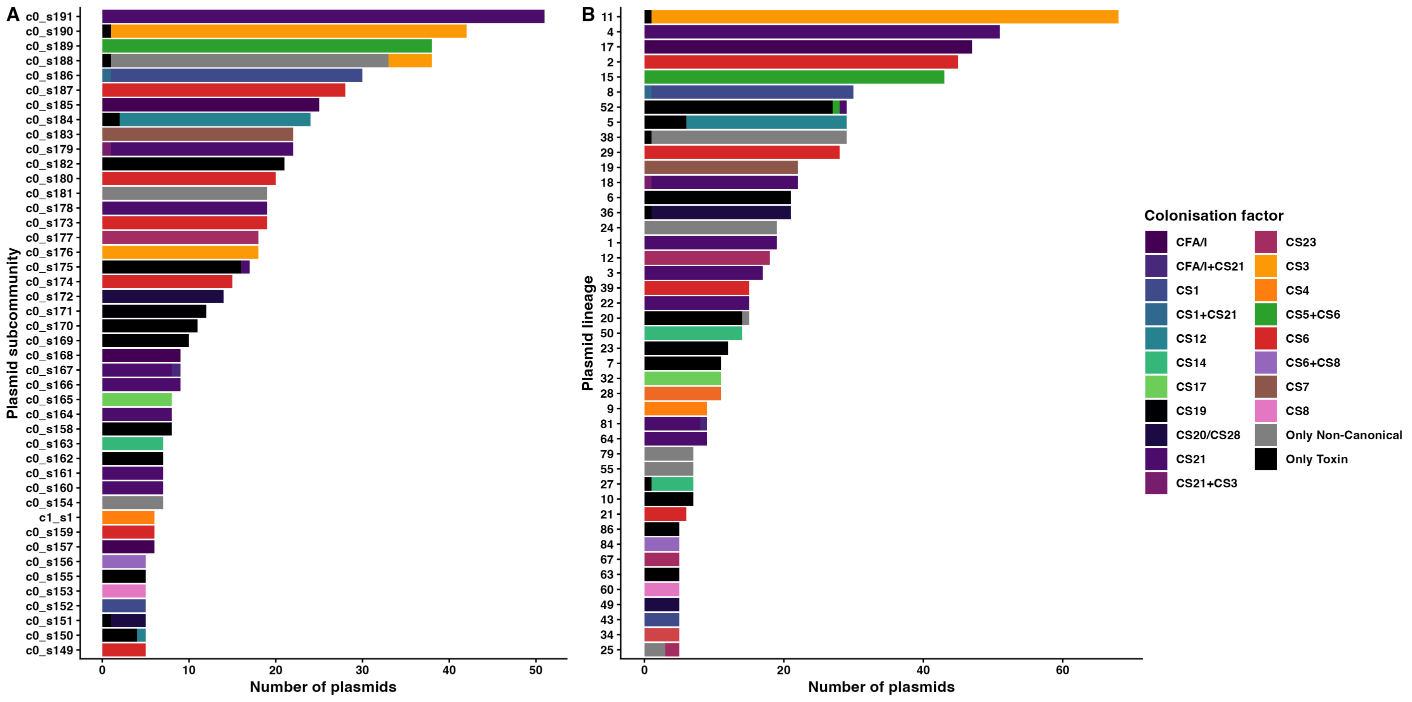


**Extended Data Fig. 3:** Pling analysis of the ETEC plasmids. (A, B): Bar charts displaying the frequency of the distinct plasmid types (Pling subcommunities) (A) and plasmid lineages (B) filled according to the combination of CFs present in the plasmid. C: community, s: subcommunity. Only groups with more than 5 isolates are shown.

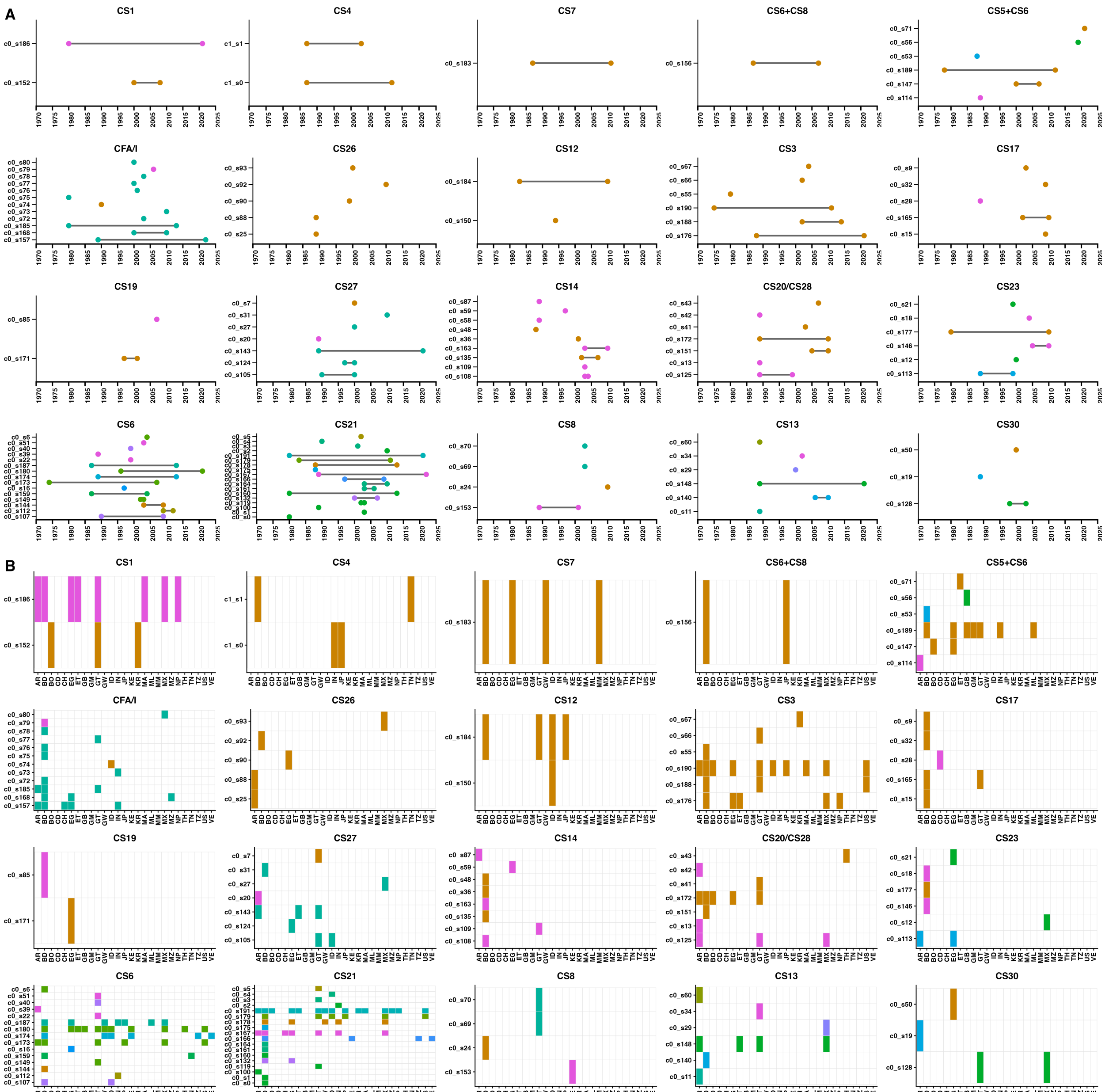


**Extended Data Fig. 4:** Plasmid maintenance through time and countries. (A) Lollipop plots showing the first and last year of detection of each plasmid subcommunity for each CF. (B): Heatmaps showing the presence of each plasmid subcommunity across different countries for each CF. (only including those with more than five isolates). Colours indicate the plasmid lineage to which each subcommunity belongs. Only CFs present in more than five isolates are shown.

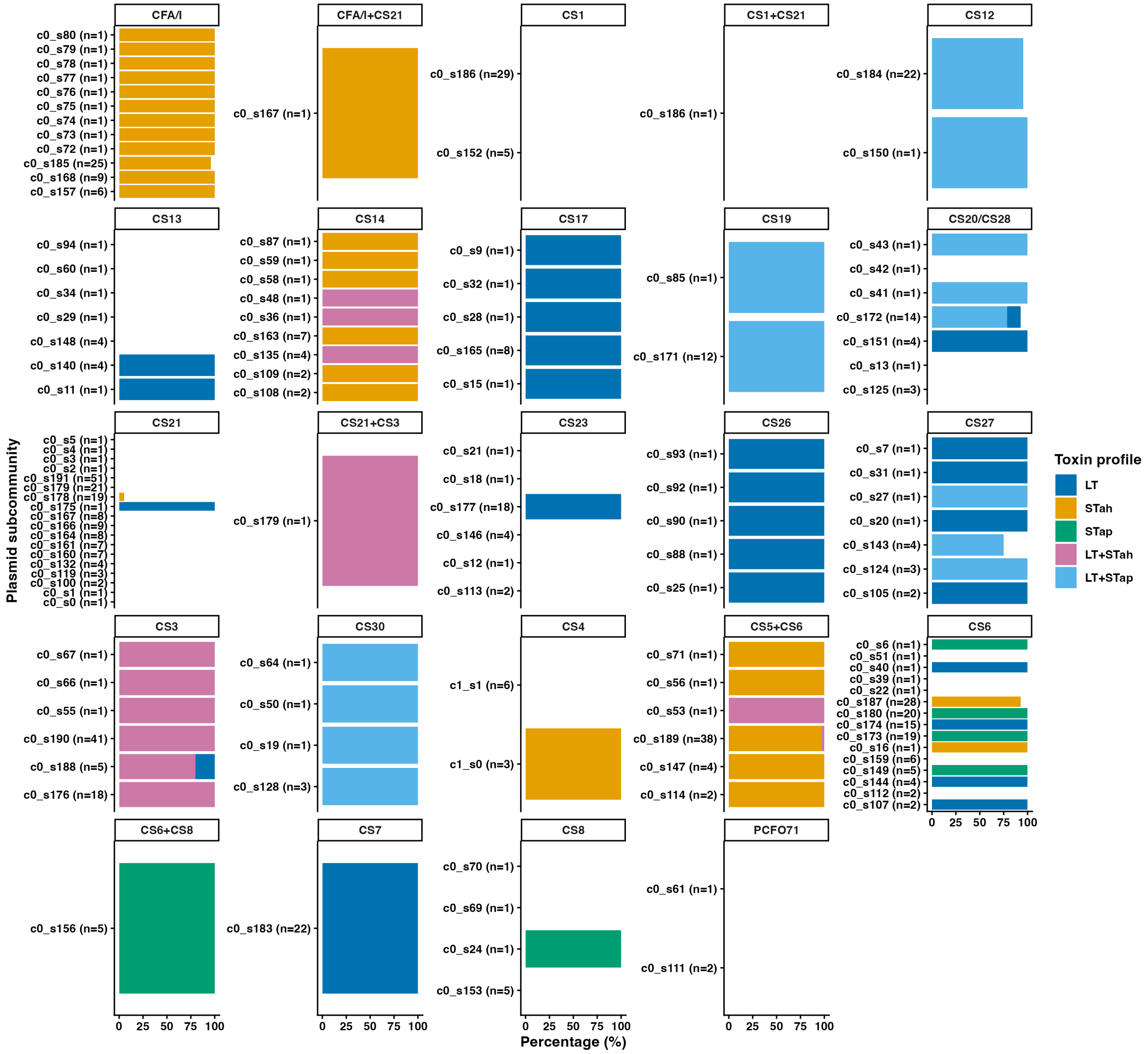


**Extended Data Fig. 5:** Bar chart displaying the frequency of each plasmid subcommunity for each combination of colonisation factors, filled by the presence of the distinct toxin profiles.

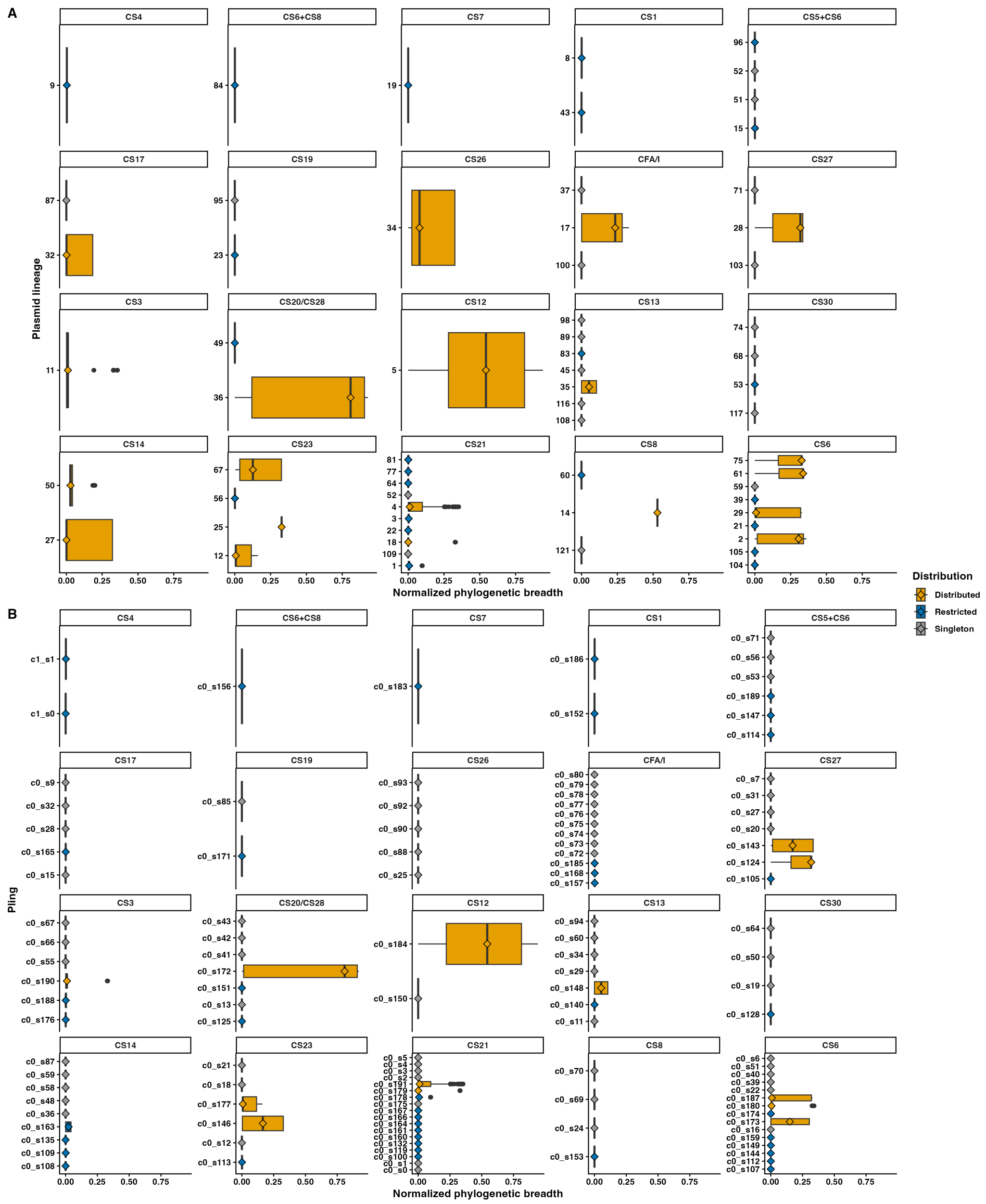


**Extended Data Fig. 6:** (A, B): Boxplots displaying the normalized phylogenetic breadth (core genome distance) for all the distinct plasmid lineages (A) and Pling subcommunities (B) for the CF (only including those with more than 5 isolates). Plasmid lineages or subcommunities were considered distributed when the maximum normalized phylogenetic breadth was greater than 0.1. Boxplots were coloured according to whether the lineages were restricted (blue), distributed (orange) or simply singletons (grey).

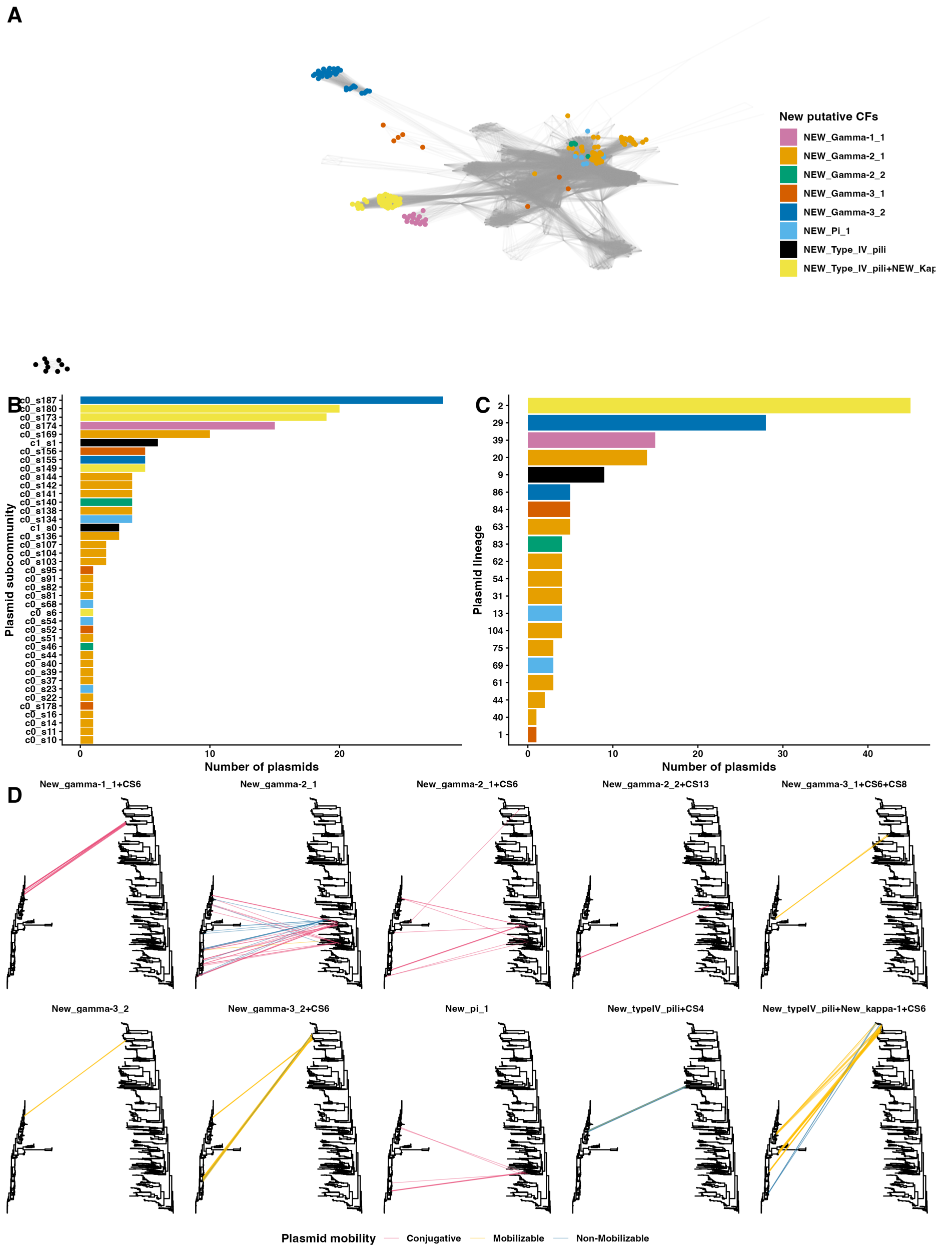


**Extended Data Fig. 7: a,** Plasmid rearrangement distance network coloring the presence of the new putative adhesive molecules. (B, C): Bar chart displaying the frequency of the distinct Pling subcommunities (B) and plasmid lineages (C) coloured according to the presence of the new putative adhesive molecules present in the plasmid. C: community, s: subcommunity. Only groups with presence of new putative adhesive molecules are shown. (D): Tanglegrams showing the distribution of the combination of putative new adhesive molecules with known CFs plasmids into core genome. Tree to the left represents a maximum-likelihood core genome phylogeny and the tree to the right represents a plasmid neighbour-joining phylogeny based on Pling containment distances. Colour of the connection lines represent the predicted plasmid mobility. Only combinations present in more than 5 plasmids are shown.

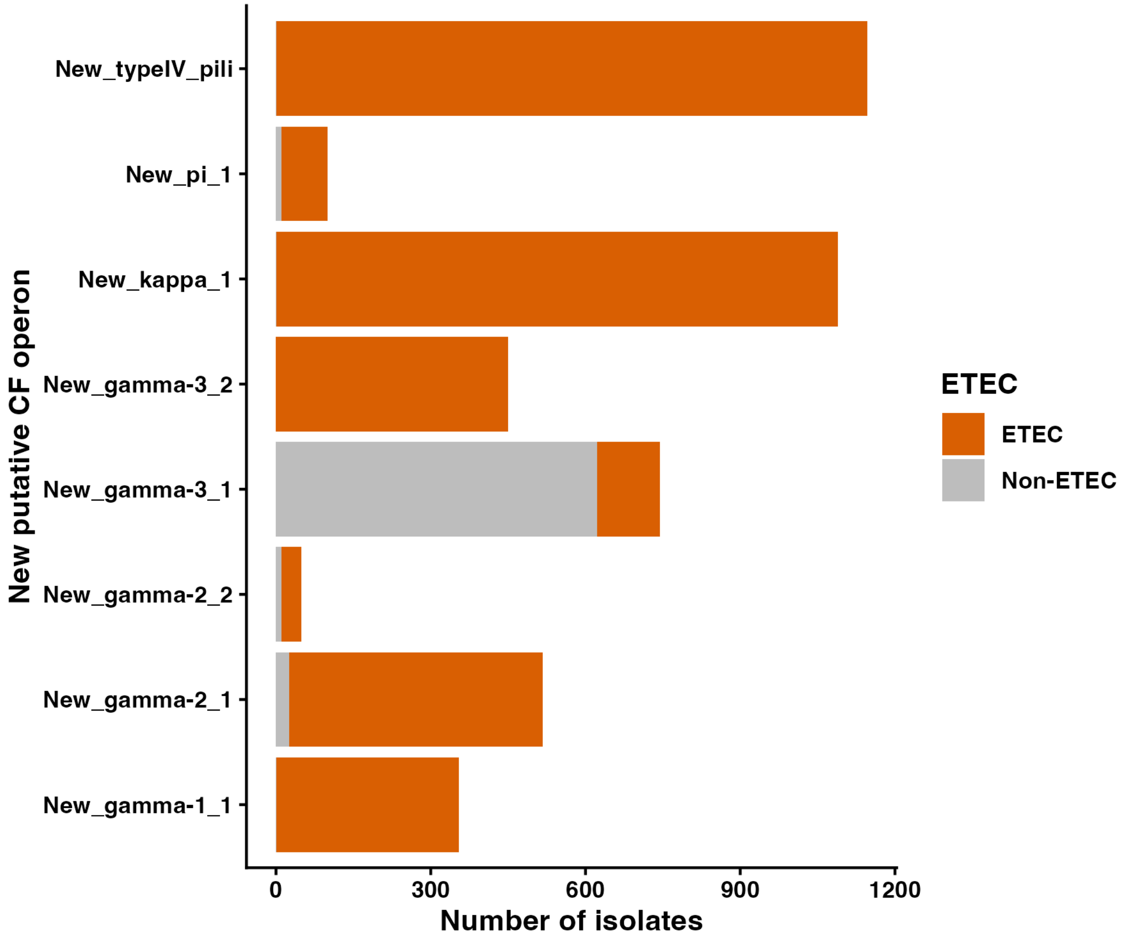


**Extended data Fig. 8:** Bar chart displaying the number of isolates in AllTheBacteria database carrying one of the new putative adhesive molecules coloured by the presence (orange) or absence (grey) of an enterotoxin.

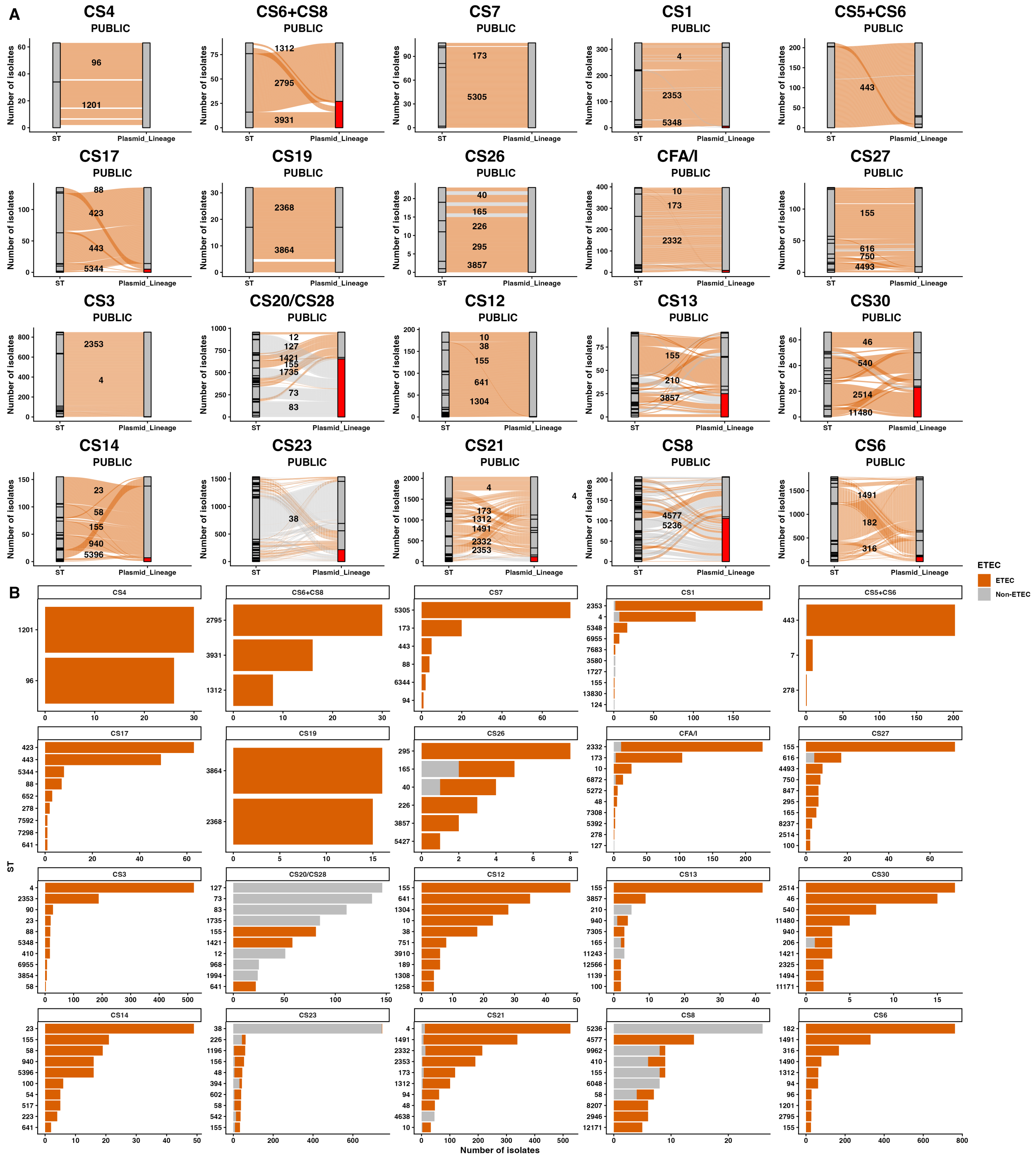


**Extended Data Fig. 9:** (A): Alluvial plots displaying the distribution of each plasmid lineage into the different STs in short-read assemblies publicly available in AllTheBacteria dataset. Left stratum represents Sequence Type (ST), and the right stratum represents Plasmid Lineage. The colour red in the right stratum represents a different plasmid from those detected in our study. For visualisation purposes, only STs present in more than 5% of the genomes containing a specific CF were labelled. (B): Bar plots displaying the 10 most common ST of a CF for known CFs present in more than 5 isolates in the dataset of this study filled by the presence or absence of an enterotoxin. Colour represents if an isolate belongs to ETEC pathotype (orange, carrying an enterotoxin) or not (grey)
